# Mutant SETBP1 enhances NRAS-driven MAPK pathway activation to promote aggressive leukemia

**DOI:** 10.1101/2021.02.01.429244

**Authors:** Sarah A. Carratt, Theodore P. Braun, Cody Coblentz, Zachary Schonrock, Rowan Callahan, Brittany M. Smith, Lauren Maloney, Amy C. Foley, Julia E. Maxson

## Abstract

Mutations in SET binding protein 1 (SETBP1) are associated with poor outcomes in myeloid leukemias. In the Ras-driven leukemia, juvenile myelomonocytic leukemia, SETBP1 mutations are enriched in relapsed disease. While some mechanisms for SETBP1-driven oncogenesis have been established, it remains unclear how SETBP1 specifically modulates the biology of Ras-driven leukemias. In this study, we found that when co-expressed with Ras pathway mutations, SETBP1 promoted oncogenic transformation of murine bone marrow in vitro and aggressive myeloid leukemia in vivo. We demonstrate that SETBP1 enhances the NRAS gene expression signature, driving upregulation of mitogen-activated protein kinase (MAPK) signaling and downregulation of differentiation pathways. SETBP1 also enhances NRAS-driven phosphorylation of MAPK proteins. Cells expressing NRAS and SETBP1 are sensitive to inhibitors of the MAPK pathway, and treatment with the MEK inhibitor trametinib conferred a survival benefit in a mouse model of NRAS/SETBP1-mutant disease. Our data demonstrate that despite driving enhanced MAPK signaling, SETBP1-mutant cells remain susceptible to trametinib in vitro and in vivo, providing encouraging pre-clinical data for the use of trametinib in SETBP1-mutant disease.

## To the Editor

SET binding protein 1 (SETBP1) mutations are associated with relapsed disease and reduced survival in juvenile myelomonocytic leukemia (JMML), a rare form of early childhood leukemia driven by Ras pathway mutations (NF1, NRAS, KRAS, PTPN11 and CBL)^1^. SETBP1 mutations occur in as many as 30% of JMML patients and reduce the five-year event-free survival rate from 51% to 18%^2^, ^3^. Although some mechanisms of oncogenesis have been established for SETBP1 mutations^4-7^, it remains unclear why they are associated with poor prognosis and relapse in this context. The goal of this study was to understand how SETBP1 modulates the biology of Ras-driven leukemias and to determine whether there are therapeutic vulnerabilities that can be exploited.

Mutations in SETBP1 are localized within its degron motif and lead to SETBP1 overexpression at the protein level^4^. SETBP1 overexpression promotes oncogenesis by binding SET, protecting the SET protein from protease cleavage^5^. Stabilization of SET leads to the inhibition of the tumor suppressor PP2A protein through the formation of a SETBP1-SET-PP2A complex^5^. SETBP1 also acts as a transcriptional regulator, and its mutation perturbs transcription of RUNX1^6^, HOXA9 and HOXA10^7^, regulators of hematopoiesis. To address our central question of how SETBP1 mutations modulate Ras-driven leukemia, we first set out to determine whether mutant SETBP1 promotes the growth of hematopoietic progenitors with a Ras pathway mutation. As there are no models of endogenous SETBP1 mutation, we leveraged retroviral vectors to express SETBP1, mimicking protein overexpression driven by the endogenous mutation. The models developed for this study can be leveraged for future drug development efforts and mechanistic studies of SETBP1 mutant disease.

SETBP1 mutations are known to co-occur with both PTPN11 and NRAS mutations (such as PTPN11^E76K^ and NRAS^G12D^)^1, 2, 8^. We find that, in the absence of exogenous cytokines, both PTPN11^E76K^ (Fig.1A, B) and NRAS^G12D^ (Fig.1C, D) formed a modest number of murine hematopoietic colony forming units (CFU). The addition of SETBP1^D868N^ significantly augmented colony number with either Ras pathway mutation. To determine whether the addition of a SETBP1 mutation enhanced self-renewal, we performed a serial replating assay with cytokine-free methylcellulose. PTPN11^E76K^ and SETBP1^D868N^ confer replating potential out to the third plating (Fig.1B). The combination of NRAS^G12D^ and SETBP1^D868N^ confer robust serial replating out to at least the fourth plating (Fig.1D), indicating that the SETBP1^D868N^ enhances both oncogenic transformation and self-renewal.

**Figure 1.**
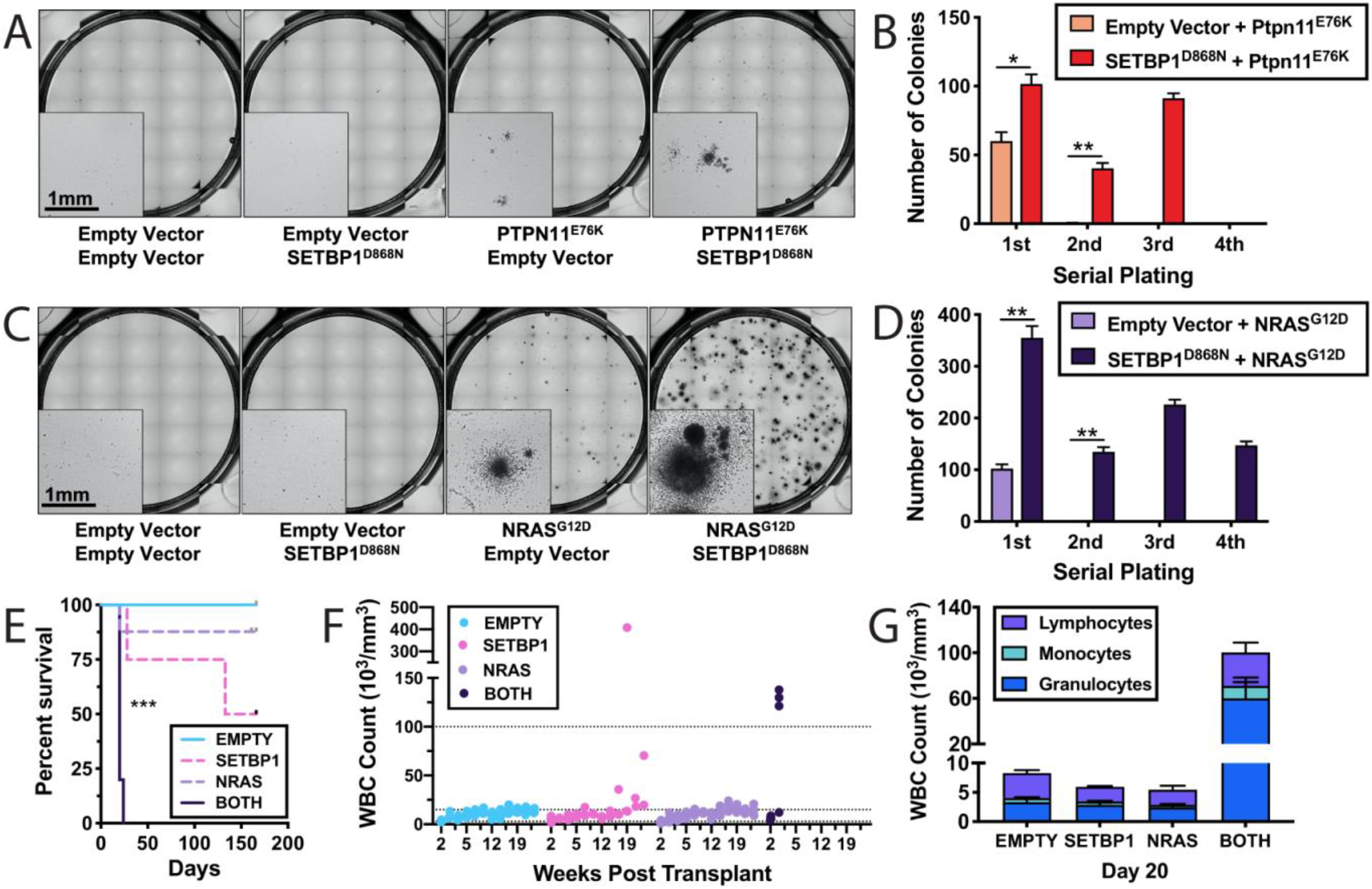
Mutant SETBP1 enhances the proliferation of NRAS-mutant hematopoietic progenitors. **(A)** Representative images of colony forming unit (CFU) assays show that SETBP1^D868N^ enhances the colony forming capacity of PTPN11^E76K^ . **(B) Quantification showing that** PTPN11^E76K^ synergizes with SETBP1^D868N^ to produce significantly more colonies than with PTPN11^E76K^ alone. Serial replating of cells co-transfected with PTPN11^E76K^ and SETBP1^D868N^. **(C)** Representative images show an enhancement of colony forming potential in cells transduced with both NRAS^G12D^ and SETBP1^D868N^ mutations. While NRAS^G12D^ alone produces some small dense colonies, the combination of NRAS^G12D^ and SETBP1^D868N^ results in an increased number of colonies that were generally larger than with NRAS^G12D^ alone. **(D)** Quantification of colony number showing that when combined with SETBP1^D868N^ mutation, the NRAS^G12D^ mutation produces significantly more colonies than with NRAS^G12D^ alone. Enhanced serial replating of NRAS^G12D^ and SETBP1^D868N^ expressing progenitors relative to those expressing NRAS^G12D^ alone. Statistical significance is represented as *p<0.05, **p<0.01. **(E)** Lineage-depleted mouse bone marrow cells expressing SETBP1^D868N^ and/or NRAS^G12D^ with the appropriate retroviral control vectors were transplanted into lethally irradiated mice with carrier bone marrow cells. The median survival in mice with BOTH oncogenes (SETBP1^D868N^/ NRAS^G12D^) was 20 days, compared to 149.5 days with SETBP1 alone (SETBP1^D868N^/Empty). Mice receiving cells expressing NRAS alone (Empty/NRAS^G12D^) did not reach their median survival by 165 days. Significance was determined by logrank (Mantel-Cox test), with the threshold for significance of p-value < 0.0083. **(F)** At 20 days post-transplant (the median survival time for mice with BOTH mutations together (SETBP1^D868N^/ NRAS^G12D^)), a marked elevation of white blood cells is seen relative to all other groups. **(G)** Peripheral complete blood counts were monitored over time. Mice expressing mice with BOTH oncogenes (SETBP1^D868N^/ NRAS^G12D^) developed high WBC counts in the first three weeks, while mice with SETBP1 only (SETBP1^D868N^/Empty) began to develop high WBC counts after 17 weeks. CBCs are shown for each group at day 20.

To understand whether mutant SETBP1 augments the oncogenicity of NRAS *in vivo*, we performed a murine bone marrow transplant experiment. For this study, 5,000 lineage-depleted murine hematopoietic cells expressing either NRAS^G12D^, SETBP1^D868N^, or both were transplanted into lethally irradiated C57BL/6J mice with 200,000 carrier cells. NRAS^G12D^/SETBP1^D868N^ mice developed an aggressive myeloid leukemia with a median survival of 20 days (Fig.1E). In contrast, the mice transplanted with SETBP1^D868N^ or NRAS^G12D^ alone, developed disease with a much longer latency (Fig.1E). The NRAS^G12D^/SETBP1^D868N^ mutant leukemia was marked by high peripheral leukocytosis (Fig.1F), an expansion of the myeloid compartment (Fig.1G, Fig.S1B), and splenomegaly (Fig.S1A).

To understand how NRAS^G12D^ and SETBP1^D868N^ cooperate to produce this aggressive phenotype, we performed RNAseq on transduced lineage-depleted hematopoietic cells. The impact of SETBP1^D868N^ alone is subtle, with only nine differentially expressed genes for SETBP1^D868N^ alone relative to control. We found that cells expressing both NRAS^G12D^ and SETBP1^D868N^ have 803 differentially expressed genes relative to the empty vector control, of which approximately half are also significant in the NRAS^G12D^ alone condition, indicating that their altered expression is primarily driven by NRAS^G12D^ (Fig.2A). We identified 399 differentially expressed genes in the NRAS^G12D^ plus SETBP1^D868N^ condition (BOTH) that are *not* differentially expressed with either oncogene alone. KEGG mouse pathway analysis of these 399 genes suggests that they are involved with MAPK activation and regulation of hematopoietic differentiation.

**Figure 2.**
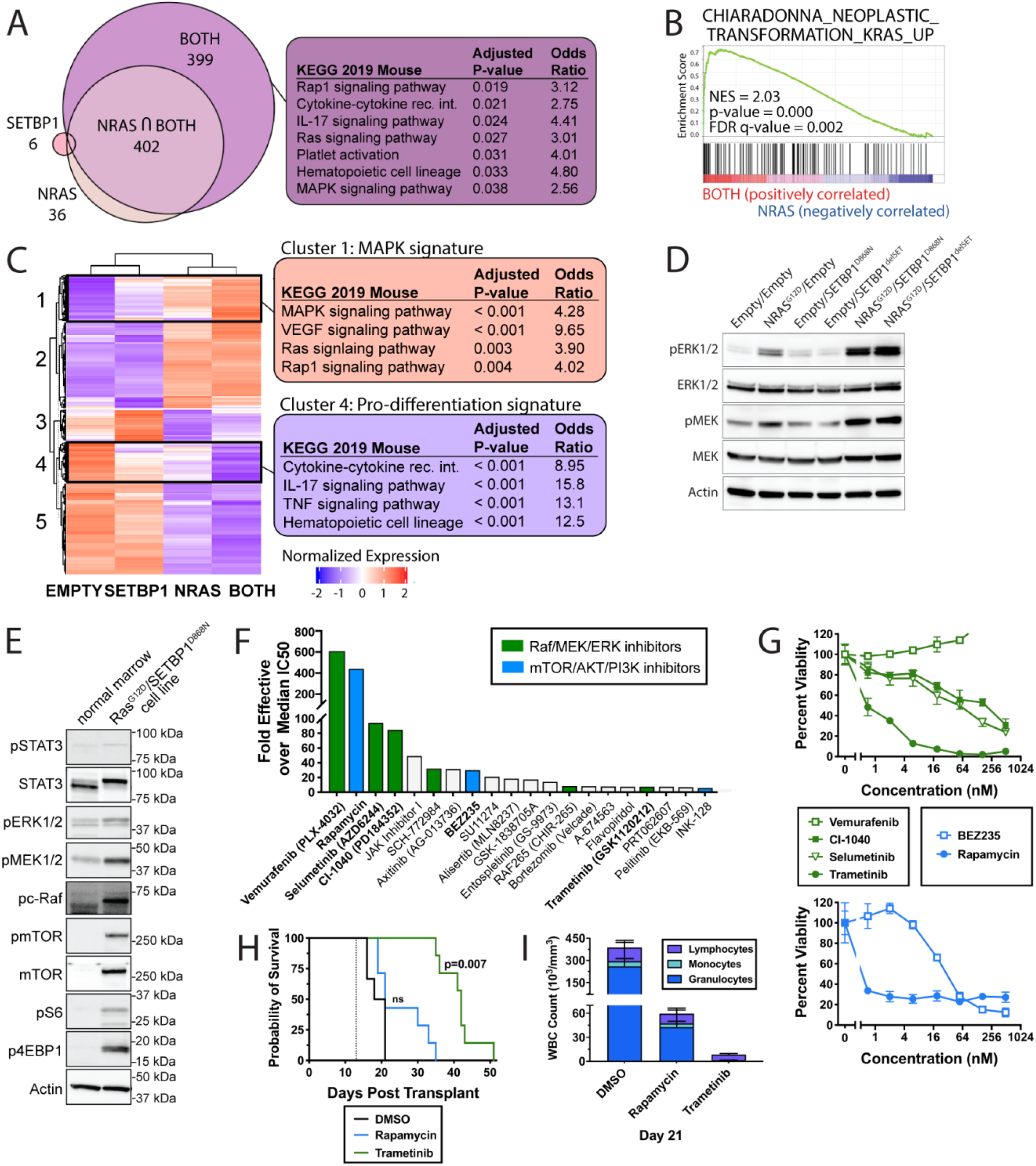
SETBP1^D868N^ enhances MAPK signaling driven by NRAS^G12D^. **(A)** RNAseq differential expression analyses were performed on transduced lineage depleted murine bone marrow cells expressing empty vector (Empty/Empty), SETBP1 (SETBP1^D868N^/Empty), NRAS (Empty/ NRAS^G12D^) or BOTH oncogenes (SETBP1^D868N^/NRAS^G12D^). The Venn diagram shows the number of genes that are differentially expressed relative to the empty vector control for SETBP1, NRAS and BOTH (logFC +/- 1.5, adj p-value < 0.05). There were 402 differentially genes at the intersection of NRAS and BOTH, and 399 genes that were differentially expressed only in the BOTH group. Enrichr analysis of these genes that are only differentially expressed with BOTH oncogenes showed upregulation of inflammatory and Ras/MAPK pathways. **(B)** Gene Set Enrichment Analysis (GSEA) was used to determine what pathways were driving differences between the NRAS-only and BOTH conditions. In the BOTH condition relative to the NRAS condition, there is an enrichment for genes upregulated by NRAS. **(C)** Unsupervised clustering of differentially-expressed genes. Cluster 1 shows a strong MAPK signature in genes upregulated by both SETBP1 and NRAS. Cluster 4 is enriched for KEGG pathways associated with myeloid differentiation. **(D)** Immunoblot analysis of 293T17 cells transiently transfected with and empty vector alone, NRAS^G12D^, SETBP1^D868N^ or the combination of both genes. Empty vector control are used to control for the total amount of plasmid transfected. Co-transfection with NRAS^G12D^ and SETBP1 increases the phosphorylation of ERK and MEK above NRAS^G12D^ alone. Deletion of the SET-binding domain from SETBP1 (SETBP1^delSET^) does not reduce MEK/ERK activation relative to full length SETBP1^D868N^. (**E**) Immunoblot analysis of NRAS^G12D^/ SETBP1^D868N^ expanded hematopoietic progenitors reveals increased activation of MAPK and mTOR signaling relative to a normal marrow control. **(F)** A chemical screen with commercially available inhibitors was performed on our novel NRAS^G12D^/SETBP1^D868N^ cell line to identify essential cell growth and survival pathways, and the cells were found to be highly sensitive to Raf/MEK/ERK inhibitors (black). **(G)** To validate the efficacy of identified inhibitors against the NRAS^G12D^/SETBP1^D868N^ cells, a 7-point dose response curve with a fixed molar ratio of each of the top agents was performed, and a percent viability calculated relative to untreated cells after 72 hours. Trametinib displayed sub-nanomolar efficacy. **(H)** To evaluate the efficacy of trametinib *in vivo*, 100,000 NRAS^G12D^/SETBP1^D868N^-mutant cells were retro-orbitally injected into C75BL/6J mice without irradiation. Beginning at day 13 (dotted vertical line), mice were given once-daily treatment of either DMSO or 1 mg/kg trametinib. Median survival in mice receiving the DMSO control treatment was 19.5-days post-transplant compared 42 days with trametinib (p=0.0007). Significance was determined by logrank (Mantel-Cox test). (**I**) At Day 21, the disease burden was markedly higher in the peripheral blood of DMSO-treated mice relative to trametinib-treated mice.

We performed Gene Set Enrichment Analysis (GSEA) to determine what pathways were enriched when SETBP1^D868N^ was co-expressed with NRAS^G12D^ compared to NRAS^G12D^ alone (Fig.2B). Surprisingly, we found that when both oncogenes are expressed, there is an enrichment for genes that are upregulated by the Ras pathway, suggesting that SETBP1 further augments the oncogenic NRAS signature. To understand this phenomenon, we performed unsupervised clustering on all genes that were differentially expressed between any of the conditions (Fig.2C). As expected, a majority of the signaling changes relative to Empty could be attributed to NRAS^G12D^ (Clusters 2, 5). However, we identified two clusters in which genes had increased or decreased expression when SETBP1^D868N^ was co-expressed with NRAS^G12D^ versus either oncogene alone (Clusters 1, 4). Cluster 4 is defined by pro-differentiation pathways that have the most downregulation with the combination. Cluster 1 is defined by genes that are upregulated to a small degree by each NRAS and SETBP1 individually, with the greatest upregulation in the combination. Cluster 1 has a strong MAPK signature (Fig.2C). To understand whether the enhanced transcriptional activation downstream of the MAPK pathway by SETBP1 was accompanied by an increase in MAPK phosphorylation and activation, 293T17 cells were transiently transfected with NRAS^G12D^ and/or SETBP1^D868N^ (Fig.2D). While SETBP1^D868N^ alone does not increase signaling in the MAPK pathway over baseline expression, in the context of a NRAS^G12D^ mutation SETBP1^D868N^ augments phosphorylation of ERK and MEK, two key MAPK pathway proteins. To understand whether SETBP1^D868N^ modulates the MAPK pathway through it’s known interacting partner, SET, we deleted the SET binding domain from SETBP1 and found no difference in MAPK phosphorylation relative to the full length SETBP1 (Fig.2D).

To facilitate further mechanistic studies, we generated a NRAS^G12D^/SETBP1^D868N^ model by culturing transduced murine hematopoietic progenitors in IMDM with 10% serum (Fig.S1C). This combination of NRAS^G12D^ and SETBP1^D868N^ mutations (but not either mutation alone) enable cells to proliferate in culture over multiple passages. Relative to normal bone marrow, these expanded hematopoietic progenitors have increased MAPK and mTOR pathway signaling (Fig.2E). To identify dependencies in SETBP1-transformed cells we performed a chemical screen on these NRAS^G12D^/SETBP1^D868N^ hematopoietic progenitors (Fig.2F). Screening of these cells against a panel of drugs targeting cell growth and survival pathways revealed a unique sensitivity to Raf/MEK/ERK and mTOR/AKT/PI3K inhibitors when compared with all other samples previously screened on the platform^9^. A subset of these inhibitors were validated individually (Fig.2G), with most efficacious of these drugs being rapamycin and trametinib. Of note, trametinib is a MEK inhibitor that is currently in clinical trial for relapsed and refractory JMML and has an IC50 of 0.42 nM in these cells (Fig.S1D). Trametinib and rapamycin were then evaluated *in vivo*, where trametinib treatment significantly increased the median survival in this model from 19.5 days with DMSO to 42 days (Fig.2H). At Day 21, when the last of the vehicle-treated mice succumbed to disease, the average peripheral WBC count in the vehicle-treated mice was 387,000/mm^3^ compared to 8,600/mm^3^ in the trametinib-treated group (Fig.2I, Fig.S1J). Rapamycin neither improved survival over DMSO or significantly reduced leukocytosis at Day 21 (Fig.S1K). Optimization of model is reported in the supplement (Fig.SE-I).

In our murine model, the combination of NRAS^G12D^ with SETBP1^D868N^ produces a far more aggressive disease than NRAS^G12D^ alone. Surprisingly, the observed *in vitro* and *in vivo* synergy appears to be driven by the modulation of just a handful of pathways. Mutant SETBP1 suppresses myeloid differentiation programs while amplifying NRAS^G12D^-driven transcriptional signatures. SETBP1 is known to inhibit activity of the tumor suppressor PP2A, a regulator of the Ras/MAPK pathway, through the stabilization of SET^5^, ^10^. Interestingly, we find that deletion of the SET binding domain does not abrogate the enhancement of MAPK activation by SETBP1 (Figure 2D). This might suggest that other protein interactions drive the increased phosphorylation of MEK/ERK, or that SETBP1-driven transcriptional changes alter the activation potential of the pathway.

Targeting the Ras pathway has been of great interest for patients with JMML^11^, where SETBP1 mutations are associated with significantly lower five-year event free survival^2^ and additional therapeutic options are needed. A clinical trial is currently underway targeting signaling downstream of Ras with the MEK/ERK inhibitor trametinib in patients with relapsed and refractory JMML (ClinicalTrials.gov Identifier: NCT03190915). It was previously unclear whether the presence of a SETBP1 mutation would alter the efficacy of trametinib. Inactivation of PP2A by the PP2A Aα R183W point mutation has been shown to drive resistance to MEK inhibitors through the potentiation of Ras signaling and ERK phosphorylation^12^. Since SETBP1 is an inhibitor of PP2A it could theoretically reduce sensitivity to MEK inhibitors. However, we find that SETBP1-mutant cells are still highly sensitive to inhibitors of the RAS/ERK/MAPK pathway. Further, trametinib doubles overall survival in our murine model of NRAS/SETBP1-mutant leukemia, providing encouraging pre-clinical data for the use of trametinib in SETBP1-mutant JMML.

## Author Contribution

*Concept and design:* Carratt, Braun, Maxson

*In vitro experiments:* Carratt, Coblentz, Schonrock, Smith, Maloney, Foley, Maxson

*In vivo experiments:* Carratt, Schonrock

*Analysis and interpretation of data:* Carratt, Braun, Callahan, Maxson

*Writing, review and revision of the manuscript:* Carratt, Braun, Coblentz, Callahan, Schonrock, Smith, Maloney, Foley, Maxson

## Acknowledgements

Research reported in this publication was supported by NCI F32CA239422 to SAC, as well as a Knight Pilot Award, American Society of Hematology Scholar Award, and Gilead Research Scholars Program in Hematology/Oncology to JEM. The authors gratefully acknowledge the support of the OHSU Flow Cytometry Shared Resource, with particular regard to operators Brianna Garcia and Dorian LaTocha.

