## Supplement for "Mutant SETBP1 enhances NRAS-driven MAPK pathway activation to promote aggressive leukemia"

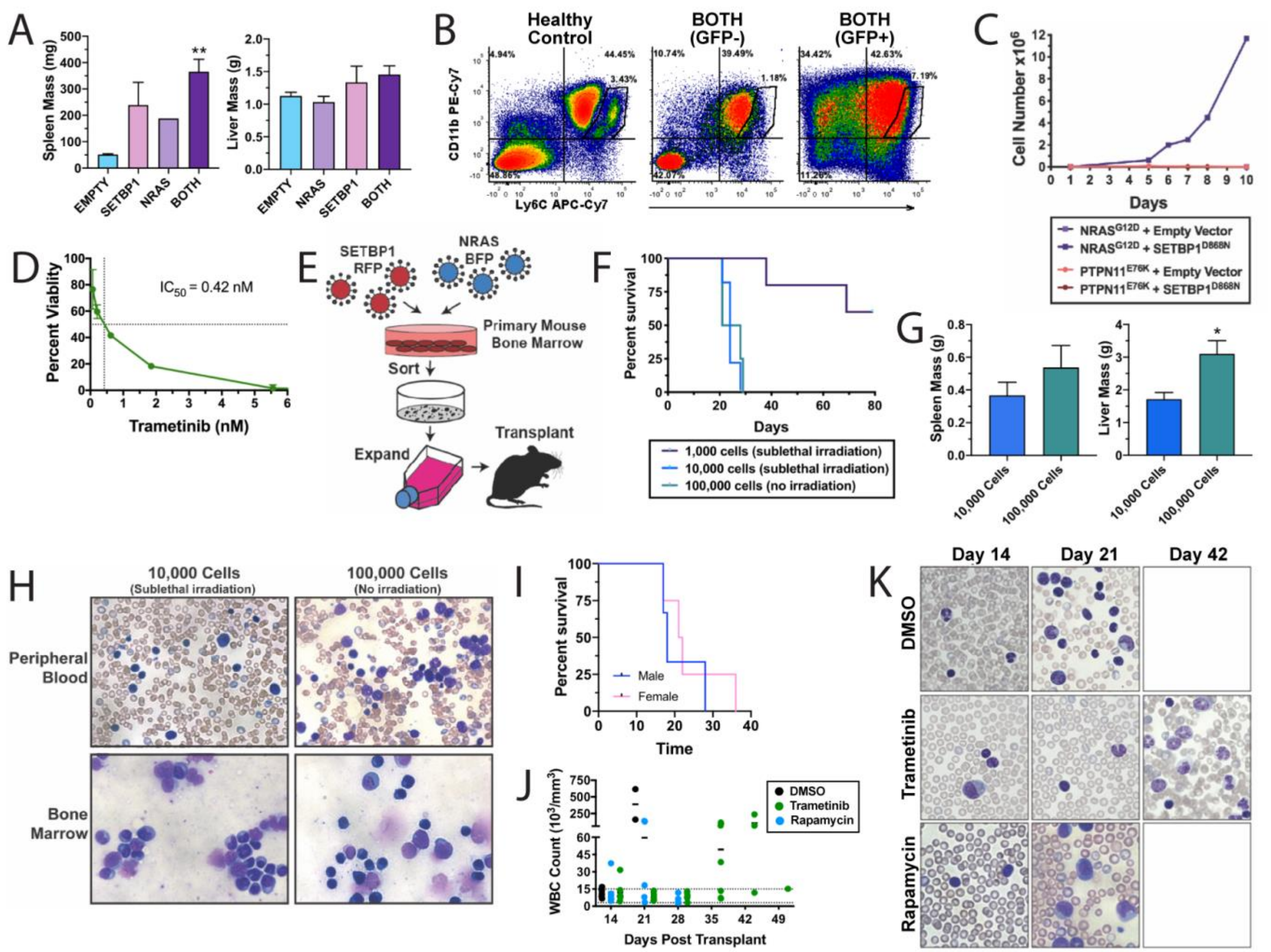

**Figure S1. (A)** Terminal spleen and liver weights from bone marrow transplant experiment with hematopoietic cells transduced with SETBP1$^{D868N}$ and/or NRAS$^{G12D}$ shown in Figure 1E-G. **(B)** Representative terminal flow cytometry in mice with BOTH mutations. Population of Ly6C$^{Hi}$/Cd11b+ cells that is present in healthy bone marrow is lost the host marrow (GFP-), indicating abnormal granulopoiesis in the presence of the NRAS+/SETBP1+ transduced marrow. The NRAS+/SETBP1+ cells (GFP+) are 77% Cd11b+ and 43%CD11b+/Ly6C+. **(C)** Mouse bone marrow cells were harvested from CFU assays (as in Figure 1A and C). Cells were washed and maintained in liquid culture (IMDM with 20% FBS) without any additional cytokine support. Within one week, only NRAS$^{G12D}$ and SETBP1$^{D868N}$ transduced cells continued to survive and proliferate, thus generating a cell line. **(D)** In order to pinpoint the IC50 of trametinib, which was lower than the concentrations evaluated in 2G, an additional dose response curve was generated. The IC50 of trametinib in our SETBP1$^{D868N}$/NRAS$^{G12D}$-mutant cell line was calculated as 0.42 nM. Cell viability was assessed in triplicate by MTS assay after 72 hours of drug treatment. **(E)** Schematic of reproducible transplant design. SETBP1$^{D868N}$/NRAS$^{G12D}$ transduced cells were harvested from colony assay and expanded to generate a cell line before being transplanted into C57BL/6 mice. Optimization of cell number, sex and irradiation is described in panels F-I. **(F)** Survival curves for mice transplanted with 1,000 or 10,000 cells with sublethal (500RAD) irradiation compared to 100,000 cells with no irradiation. **(G)** Terminal spleen and liver weights for mice that received 10,000 cells with sublethal irradiation or 100,000 cells with no irradiation. **(H)** Peripheral blood and bone marrow smears for mice that received 10,000 cells with sublethal irradiation or 100,000 cells with no irradiation. **(I)** Male and female mice transplanted with 100,000 cells and no irradiation had similar survival times. **(J)** WBC count over time as measured by automated CBC for the in vivo drug study described in Figure 2H-I. **(K)** Representative blood smears for the in vivo drug study described in Figure 2H-I. Only mice treated with trametinib survived to 42 days, at which time the surviving mice did succumb to myeloid leukemia.

**Supplemental methods**

*Mice*. C57BL/6J mice (#000664) and Balb/cJ mice (#000651), were obtained from The Jackson Laboratories. Female mice were used between 6 and 8 weeks, and age and weight matched in all experiments. All experiments were conducted in accordance with the National Institutes of Health Guide for the Care and Use of Laboratory Animals and approved by the Institutional Animal Care and Use Committee of Oregon Health & Science University (Protocol #TR01_IP00000482). Peripheral blood collected from the saphenous vein was monitored weekly for white blood cell counts (WBC) using a Vet ABC animal blood counter (Scil animal care company). The health of the mice was monitored daily by animal care personnel who were blinded to study groups. Animals were euthanized if WBC reached above 100,000/mm$^3$.

*Selection of genetic variants.* Patients with SETBP1 mutations most commonly have somatic variants within a four amino acid segment of the SKI-homology domain (1). These mutations increase the half-life of SETBP1 by perturbing the β-TrCP degron motif, such that E3 ubiquitin ligases cannot recognize the domain to degrade the protein (1). Mutations in the GTPase NRAS (such as the G12D mutation) are common in JMML and usually occur in the $Mg^{2+}$ nucleotide binding domains (2,3). Mutations in tyrosine-protein phosphatase non-receptor type II (PTPN11), a regulator of the Ras pathway, are also found in combination with SETBP1 mutations in JMML (3). For this study, we selected representative pathway mutations based on their co-occurrence in JMML (2,4): SETBP1$^{D868N}$, NRAS$^{G12D}$ and PTPN11$^{E76K}$.

*Cloning.* For these studies, a gateway pENTR vector for V5-tagged-SETBP1 with codon-optimization was synthesized commercially (Invitrogen/GeneArt). SETBP1 was mutagenized to create the D868N mutation using the Quikchange II XL Site Directed Mutatgenesis Kit (Agilent Genomics) with primers: F-gtgccgatgccggaattgctagggattgtttcc, R-ggaaacaatccctagcaattccggcatcggcac. The SET binding domain was deleted from the V5-SETBP1 construct using the primers: F-cagcaacgacaagtggccaagtctgtctcccc, R- ggggagacagacttggccacttgtcgttgctg. NRAS$^{G12D}$ (pDonor-255) was purchased from Addgene (Hs.NRAS G12D gifted by Dominic Esposito; Addgene plasmid #83176; http://n2t.net/addgene:83176; RRID:Addgene_83176). PTPN11 MSCV-IRES GFP constructs were a gift from Dr. Jeffrey Tyner. SETBP1$^{D868N}$, NRAS$^{G12D}$, PTPN11$^{E76K}$ were transferred into a Gateway converted version of pMSCV-IRES-GFP (a gift from Tannishtha Reya; Addgene plasmid #20672; http://n2t.net/addgene:20672; RRID:Addgene_20672) or pMSCV-IRES-mCherry RFP (a gift from Dario Vignali; Addgene plasmid # 52114 ; http://n2t.net/addgene:52114 ; RRID:Addgene_52114) using Gateway LR Clonase II kit (Invitrogen). Plasmid sequences were confirmed via Sanger sequencing (Eurofins Genomics).

*Murine retrovirus and transduction.* The human embryonic kidney 293T17 cell line (ATCC) was grown in DMEM (Gibco) medium with 10% fetal bovine serum (FBS, HyClone), Glutamax (Gibco), and penicillin/streptomycin (Gibco). Retrovirus was generated by co-transfecting SETBP1 and NRAS or PTPN11 plasmids and the EcoPac plasmid (provided by Dr. Rick Van Etten) into the 293T17 cells using FuGENE 6 (Promega). Viral supernatants were harvested 48 and 72 hours later. Marrow was cultured overnight in the presence of SCF, IL6, and IL3. Mouse bone marrow ($1\times10^6$ cells/well) was spinnoculated with viral supernatant, HEPES buffer, and polybrene on two subsequent days. For the spinnoculation, cells were spun at 2,500 rpm for 90 minutes at 30°C (brake turned off).

*Murine hematopoietic colony forming unit (CFU) assays*. CFU assays were performed as described previously[13]. Retrovirus was generated as described above for SETBP1, NRAS, PTPN11 or control vector and used to transduce murine bone marrow cells isolated from 6 to 10-week old female C57BL/6J mice, and cells positive for both genes were sorted on a BD FACSAria III sorter into MethoCult M3234 methylcellulose medium (StemCell Technologies). Cells were imaged using STEMvision (StemCell Technologies), blinded, and then manually counted using ImageJ (NIH). For each replating, cells were isolated from methylcellulose, washed with PBS, and then plated in triplicate.

*In vivo oncogene synergy.* Bone marrow was harvested from 7-week-old female Balb/c mice and the mature hematopoietic cells were depleted using a Miltenyi Biotec Mouse Direct Lineage Cell Depletion Kit (130-090-858) with MS Columns (130-042-201) and 30µm Pre-Separation Filters (130-041-407). Retroviral-conditioned media was produced using 293T17 cells transduced with packaging plasmid and the appropriate transfer plasmid, as described above. Sorted double-positive cells were spun down and resuspended in PBS with non-transduced, fresh carrier bone marrow. Lethally irradiated Balb/c mice (2x4.5Gy) were transplanted with 5,000 transduced RFP+/GFP+ cells and 200,000 carrier bone marrow cells per mouse, via retro-orbital injection. Groups were randomly assigned. Mice were maintained on antibiotic water for two weeks (Polymyxin B sulfate salt, Sigma-Aldrich P1004; Neomycin trisulfate salt hydrate, Sigma-Aldrich N1876).

*RNAseq*. Cells were prepared as described for the murine transplant. 15,000 cells per replicate were sorted directly into Qiagen RLT lysis buffer using a BD FACSAria Fusion cell sorter. RNA was extracted using the RNeasy micro kit (Qiagen). cDNA libraries were constructed using the Takara SmartSeq for Ultra Low Input kit and sequenced using a HiSeq 2500 Sequencer (Illumina) 100 bp SR. Raw reads were trimmed with Trimmomatic[14] and aligned with STAR[15]. Differential expression analysis was performed using DESeq2[16]. Raw p values were adjusted for multiple comparisons using the Benjamini–Hochberg method. Enrichr[17, 18] and GSEA[19, 20] were used to assess gene set enrichment for pairwise comparisons (logFC +/- 1.5).

*Western blotting*. 293T17 cells were transfected as described for colony assays, without the addition of the EcoPac packaging plasmid. At 48 hours post-transfection, cells were lysed in cell lysis buffer (Cell Signaling Technologies) containing complete mini protease inhibitor tablets (Roche). To pellet cellular debris, the lysates were spun at 12,000 r.p.m., 4 °C for 10 min, and subsequently mixed with 3× SDS sample buffer (75 mmol/L Tris (pH 6.8), 3% SDS, 15% glycerol, 8% β-mercaptoethanol, and 0.1% bromophenol blue). Samples were incubated at 95 °C for 5 min and run on Criterion 4 to 15% Tris-HCl gradient gels (Bio-Rad). Gels were transferred to PVDF membranes and blocked in Tris-buffered saline with 0.05% Tween (TBST) with 5% bovine serum albumin (BSA). Horseradish peroxidase conjugated secondary antibodies against mouse IgG and rabbit IgG (CS) were used followed by imaging of the blots on a BioRad ChemiDoc™ Imaging System. Antibodies are listed in the Key Resources Table (Supplementary Table 1). Immunoblot images were captured using a BioRad ChemiDoc™ Imaging System and cropped in BioRad Image Lab Version 5.2.1 build 11.

*Flow cytometry.* Bone marrow from the femur of mice with terminal disease was lysed with SCK lysing buffer and washed. One million cells were resuspended in FACs buffer and stained with Cd11b PE-Cy7 (BD 552850, clone M1/70) and Ly6C APC-Cy7 (Biolegend 128026, clone HK1.4) for one hour. Cells were analyzed using a BD FACSAria III and FSC Express 7 Research software.

*Cell line generation and small molecule inhibitor screening.* Transduced mouse bone marrow cells harvested from CFU assays were washed four times with phosphate buffered saline (PBS, Gibco) and resuspended in IMDM (Gibco) with 20% FBS, and penicillin/streptomycin. Total cell number was monitored regularly until growth was exponential without any additional cytokine support (data not shown). A mid-throughput chemical screen was performed as described previously[21], on the NRAS$^{G12D}$/SETBP1$^{D868N}$-mutant cell line to identify dependencies. An IC50 was calculated for each inhibitor based on the 7-point dose response curve (10-0.014 nM). This IC50 was expressed relative to the median IC50 for previously screened cell lines and human samples, thus allowing for generation of a measure of the fold efficacy[21].

*In vivo trametinib efficacy.* 100,000 cells from our SETBP1$^{D868N}$/NRAS$^{G12D}$ cell line were retro-orbitally injected into C57BL/6J mice (N=6-7) with no irradiation. Mice were monitored for onset of disease by investigators and blinded animal care personnel, who monitored weight and WBC counts. At Day 13, mice were randomly assigned to treatment groups by a blinded third party and were given once-daily treatment of either 1 mg/kg trametinib (N=7) or an equivalent volume of DMSO in vehicle (N=6). Trametinib (GSK1120212) was purchased from Selleck Chemicals, resuspended in DMSO and kept in aliquots at 4 degrees until time of oral gavage, when it was then diluted in 1% sodium carboxymethyl cellulose (100 μl/dose). All mice received 8.8% dimethyl sulfoxide (DMSO) by volume.

*Data analysis and presentation.* All graphs were made using GraphPad Prism; figures were assembled in Adobe Illustrator. Western blot images were straightened and cropped in BioRad Image Lab Version 5.2.1 build 11. Data is presented as mean ± standard error of the mean. For oncogene synergy CFU assays, a nonparametric Welsh's T-test with Bonferroni posthoc correction for multiple comparisons was performed to determine significance at each timepoint. For the trametinib-treated CFU analysis, a two-way ANOVA with Dunnett corrections for multiple comparisions was used to compare each treatment to the untreated controls. For survival analysis, a logrank Mantel-Cox test was performed with Bonferroni posthoc correction for multiple comparison. In Figure 3, the Venn Diagram was generated using eulerr (R package v6.1.0) and the heatmap was created using ComplexHeatmap (v2.2.0), RColorBrewer (v1.1-2), clusterProfiler (v3.14.3), tidyverse (v1.3.0), and dplyr (v1.0.2).

*Data sharing and key resources.* Contact Julia Maxson at for information regarding renewable materials, datasets, and protocols. Raw sequencing files were deposited to the GEO repository: GSE158379. Key resources and Research Resource Identifiers (#RRID) are provided in Supplementary Tables 1 and 2.

**Supplementary Table 1. Key Resources Table**

| Reagent or Resource | Resource Type | Source | Product Number/Identifier | Dilution | Lot Number |
|---|---|---|---|---|---|
| phosphoERK1/2 | Antibody | Cell Signaling Technologies | Cat# 4370, RRID:AB_2315112 | 1:5000 | Lot 24 |
| ERK1/2 #4695 | Antibody | Cell Signaling Technologies | Cat# 4695, RRID:AB_390779 | 1:5000 | Lot 28 |
| phosphoMEK | Antibody | Cell Signaling Technologies | Cat# 9126, RRID:AB_331778 | 1:5000 | Lot 3 |
| MEK | Antibody | Cell Signaling Technologies | Cat# 9154, RRID:AB_2138017 | 1:5000 | Lot 22 |
| beta-Actin | Antibody | Cell Signaling Technologies | Cat# 4970, RRID:AB_2223172 | 1:5000 | Lot 6 |
| anti-mouse IgG, HRP linked | Antibody | Cell Signaling Technologies | Cat# 7076, RRID:AB_330924) | 1:1000 | Lot 32 |
| anti-rabbit IgG, HRP linked | Antibody | Cell Signaling Technologies | Cat# 7074, RRID:AB_2099233 | 1:1000 | Lot 26 |
| phosphoPP2A | Antibody | abcam | ab32104 | 1:500 | |
| V5 | Antibody | Invitrogen | P/N 46-0705 | 1:5000 | Lot 2106326 |
| Cd11b PE-Cy7 | FACS Antibody | BD | 552850, clone M1/70 | | |
| Ly6C APC-Cy7 | FACS Antibody | Biolegend | 128026, clone HK1.4 | | |
| 293T/17 [HEK 293T/17] cells | Cell line | ATCC | RRID:CVCL_1926 | | |
| C57BL/6J mice | Mouse | The Jackson Laboratories | #000664, RRID:IMSR_JAX:000664 | | |
| Balb/cJ mice | Mouse | The Jackson Laboratories | #000651, RRID:IMSR_JAX:000651 | | |
| V5-taggedSETBP1 with codonoptimization | Plasmid | Invitrogen/GeneArt | Sequence available on request | | |
| Hs.NRAS G12D | Plasmid | Addgene, gifted by Dominic Esposito | RRID:Addgene_83176 | | |
| PTPN11 MSCV-IRES GFP | Plasmid | Dr. Jeffrey Tyner | | | |
| pMSCV-IRES GFP | Plasmid | Addgene, a gift from Tannishtha Reya | RRID:Addgene_20672 | | |

| | | | |
|---|---|---|---|
| pMSCV-IRESmCherry RFP | Plasmid | Addgene, a gift from Dario Vignali | RRID:Addgene_52114 |
| SETBP1 WT to D868N | Mutagenesis primer | Eurofins | F-gtgccgatgccggaattgctagggattgtttcc, R-ggaaacaatccctagcaattccggcatcggcac |
| SET binding domain deletion | Mutagenesis primer | Eurofins | F-cagcaacgacaagtggccaagtctgtctcccc, R-ggggagacagacttggccacttgtcgttgctg |

**Supplementary Table 2. Software and Algorithms**

| **Name** | **Identifier** |
|---|---|
| ImageJ | RRID:SCR_003070 |
| Trimmomatic | RRID:SCR_011848 |
| STAR | RRID:SCR_015899 |
| DESeq2 | RRID:SCR_000154 |
| GraphPad Prism | RRID:SCR_002798) |
| Adobe Illustrator | RRID:SCR_010279 |
| ComplexHeatmap | RRID:SCR_017270 |
| clusterProfiler | RRID:SCR_016884 |
| Enrichr | RRID:SCR_001575 |
| Gene Set Enrichment Analysis (GSEA) | RRID:SCR_003199 |
